# Measuring complex phenotypes: A flexible high-throughput design for micro-respirometry

**DOI:** 10.1101/2020.03.16.993550

**Authors:** Amanda N. DeLiberto, Melissa K. Drown, Marjorie F. Oleksiak, Douglas L. Crawford

**Author notes:** Corresponding author: (AD).

## Abstract

Variation in tissue-specific metabolism between species and among individuals is thought to be adaptively important; however, understanding this evolutionary relationship requires reliably measuring this trait in many individuals. In most higher organisms, tissue specificity is important because different organs (heart, brain, liver, muscle) have unique ecologically adaptive roles. Current technology and methodology for measuring tissue-specific metabolism is costly and limited by throughput capacity and efficiency. Presented here is the design for a flexible and cost-effective high-throughput micro-respirometer (HTMR) optimized to measure small biological samples. To verify precision and accuracy, substrate specific metabolism was measured in heart ventricles isolated from a small teleost, *Fundulus heteroclitus*, and in yeast (*Saccharomyces cerevisiae*). Within the system, results were reproducible between chambers and over time with both teleost hearts and yeast. Additionally, metabolic rates and allometric scaling relationships in *Fundulus* agree with previously published data measured with lower-throughput equipment. This design reduces cost, but still provides an accurate measure of metabolism in small biological samples. This will allow for high-throughput measurement of tissue metabolism that can enhance understanding of the adaptive importance of complex metabolic traits.

## Introduction

Understanding evolution and ecological adaptation can be enhanced by combining genomics with quantitative analyses of complex phenotypic traits [1]. This integrative approach requires sufficient sample size (i.e. 100s to 1000s) with precise measure of phenotypes, however it can be challenging to obtain economical equipment for such high-throughput quantification. To address this challenge, we present an inexpensive custom design to measure metabolism in small biological samples such as cell suspensions, individual tissues or possibly small organisms. Metabolism is a complex trait intricated in most physiological processes and is important to organismal success. Thus, metabolism is ecologically and evolutionarily important [2–11]. The effect of the environment on metabolism as well as tissue-specific variation can vary considerably among individuals and populations [12–18]. These and other data suggest that measuring metabolism can provide insights into the ecology and evolution of organisms [5].

Metabolism is typically quantified via oxygen consumption rates (MO_2_). Numerous systems to measure MO_2_ are available from companies including Unisense, PreSens, and Loligo, but each has limitations with respect to technical design, throughput capacity, and cost. For small biological samples, systems often have limited capacity (e.g. Oxygraph 2-K, OROBOS INSTRUMENTS, Innsbruck, Austria) or require expensive reagents and disposables (e.g., Seahorse XF Analyzers, Agilent, Santa Clara, CA). Therefore, these systems are not ideal for high-throughput experimental designs as it becomes time consuming and expensive to measure many samples. There is a need in the field for a simple design that can measure multiple sample simultaneously at a reduced cost. Here we present a design for a high-throughput micro-respirometer (HTMR) that increases throughput of tissue-specific metabolism while minimizing costs maintaining and maintaining efficacy. We validate the precision and accuracy of this system by measuring both *Saccharomyces cerevisiae* and substrate specific metabolism in *Fundulus heteroclitus* heart ventricles.

## Materials and methods

### Instrumentation

The HTMR consists of a custom external plexiglass water bath designed to enclose 1-ml micro-respiration chambers (Unisense) (Fig 1A). The water bath is connected to a temperature-controlled, re-circulating system, and placed on a multi-place stir plate. Each chamber contains a stir bar and nylon mesh screen for mixing media while keeping tissues suspended (Fig 1B). Exact chamber volumes were determined by measuring the mass (to 0.001 g) of water that completely filled individual chambers with the mesh screen and stir bar. A fluorometric oxygen sensor spot (PreSens) is adhered to the internal side of the chamber lid with a polymer optical fiber cable affixed to each chamber lid for contactless oxygen measurement through the sensor spot. All cables are connected to a 10-channel microfiber-optic oxygen meter (PreSens), which uses PreSens Measurement Studio 2 software to collect oxygen data at a sampling rate of 20 measurements per minute. Sensors were calibrated at 0% (using 0.05 g sodium dithionite per 1 ml of media) and 100% air saturation (fully oxygenated media). Validation of the HTMR was carried out using a four-chamber system; however, it can easily be extended to a 10-chamber system as in Fig 1A.

**Fig 1.**
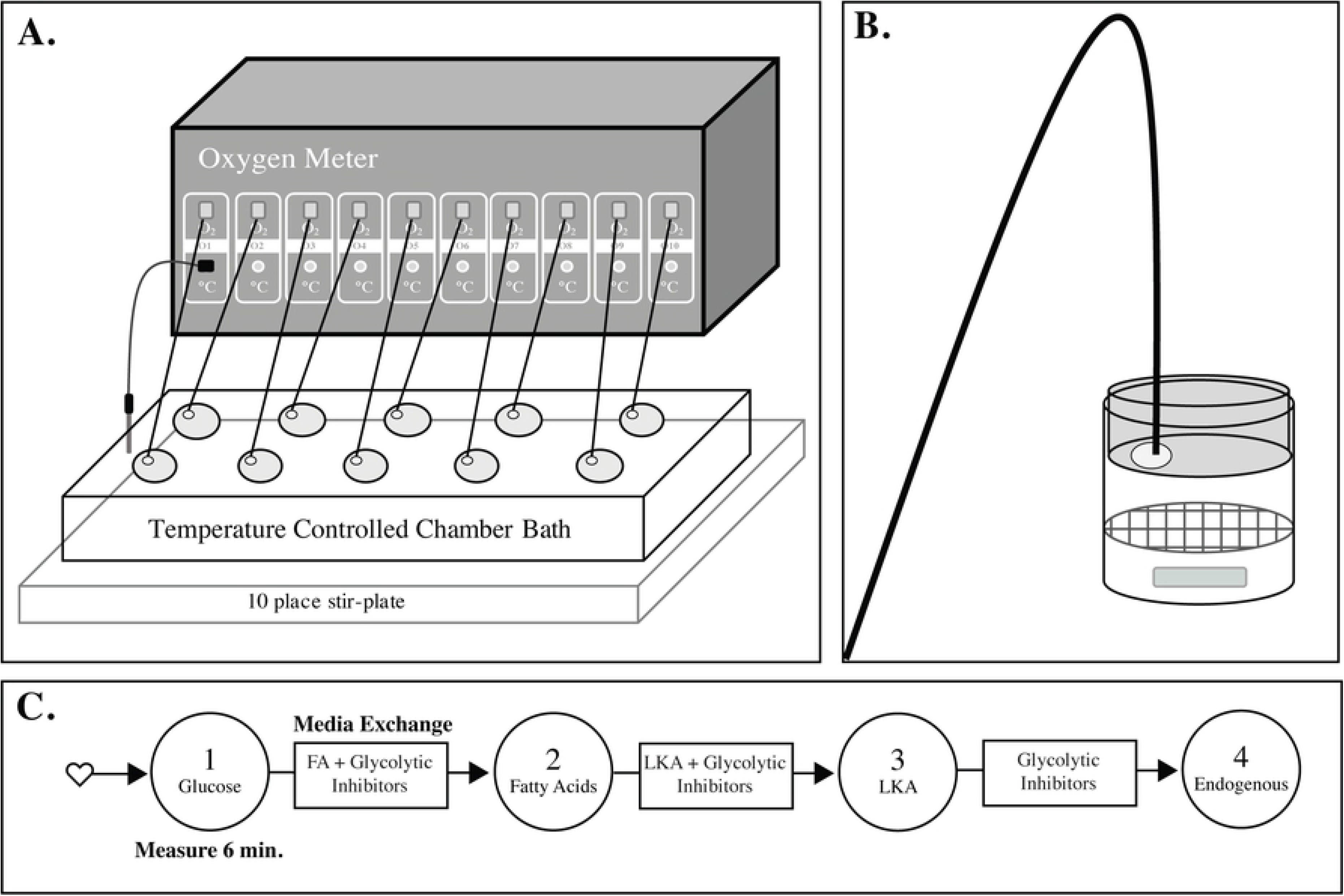
Ten chamber high-throughput micro-respirometer (HTMR). (A) PreSens 10-channel oxygen meter with fiber optic cables connected via contactless oxygen sensor spots in each chamber. Glass chambers are sealed in a water bath for constant temperature control, positioned on a stir-plate to circulate the media within each individual chamber. A temperature probe is connected and placed in the water bath to monitor temperature. (B) Individual chamber design, with Unisense 1 ml micro-respiration chamber, modified with a contactless sensor spot adhered to the inner surface of the lid and a stir bar below a nylon mesh screen. The fiberoptic cable is attached to the lid adjacent to the sensor spot. (C) General schematic of experimental metabolic measurements in teleost ventricles. Each ventricle is measured in each substrate for six minutes and then placed in the following substrate while exchanging chamber media. Substrate conditions are measured as follows: 1) 5 mM glucose; 2) 1 mM palmitic acid; 3) 5 mM lactate, 5 mM hydroxybutyrate, 5 mM acetoacetate, and 0.1% ethanol; and 4) endogenous metabolism with no substrate. Glycolytic enzyme inhibitors (10 mM iodoacetate and 20 mM 2-deoxyglucose) are added to fatty acid, LKA, and endogenous conditions to inhibit residual glycolytic metabolism.

### Metabolic rate determinations

The precision and accuracy of the HTMR was validated by measuring MO_2_ in both yeast (*Saccharomyces cerevisiae)* and teleost *(F. heteroclitus)* heart ventricles. MO_2_ is measured in the sealed chambers by measuring oxygen concentration at a rate of 20 measurements per minute, over a six-minute period. During each daily measurement, a minimum of three blank measurements, during which only media was in the chamber, were run to determine any background flux. For each six-minute measurement, the last three minutes (60 datapoints) were used for calculating metabolic rate. To do so, oxygen concentration was regressed against time to determine the raw oxygen consumption rate (pmol*μl-^1^*min^−1^). Slopes were also calculated for each blank measurement and averaged by chamber to quantify background flux, then subtracted from each slope. Metabolic rate was measured as MO_2_ = (M_sample_ – M_blank_) * V_chamber_ * 1/60, where MO_2_ is the final metabolic rate in pmol*s^−1^, M is the slope of oxygen consumption per sample in pmol*μl^−1^*min^−1^, and V is the volume of each chamber in μl.

PreSens datafiles provide data per sensor with oxygen concentration (μmol*L^−1^) at each time point (minutes). An R markdown file detailing this analysis of the raw PreSens data files can be found at https://github.com/ADeLiberto/FundulusGenomics.git.

### Experimental Organisms

*Fundulus heteroclitus* were collected using wire minnow traps from Cape Cod, MA, Mt. Desert Island, ME, and Deer Isle, ME. Individuals were trapped on public land, and no permit was needed to catch these marine minnows for non-commercial purposes. All fish were common gardened at 20°C and salinity of 15 ppt in re-circulating aquaria for at least five months and then acclimated to 12°C or 28°C for at least two months prior to metabolic measurements. Fish were randomly selected, weighed, and then sacrificed by cervical dislocation. Heart ventricles were isolated and immediately placed in Ringer’s media (1.5mM CaCl_2_, 10 mM Tris-HCl pH 7.5, 150 mM NaCl, 5mM KCl, 1.5mM MgSO_4_) supplemented with 5 mM glucose and 10 U/ml heparin to expel blood. Media was incubated at the measurement temperature prior to use. Ventricles were then splayed following precedent of previous cardiac metabolism measurements in *F. heteroclitus* [19]. Splaying the hearts decreases variation and increases overall oxygen consumption rates, as greater internal surface area is exposed to the substrate media [20]. After splaying, hearts were not further stimulated, as mechanical disruption or homogenization can increase variability in oxygen consumption rates [15]. All animal husbandry and experimental procedures were approved through the University of Miami Institutional Animal Care and Use Committee (Protocol # 19-045).

### Methodological validation

In order to validate the HTMR performance, several parameters were tested: 1) net flux at multiple oxygen concentrations, 2) between-chamber variability in MO_2_, and 3) consistency of MO_2_ over time. To quantify net flux and confirm equal rates between chambers, flux was measured at multiple oxygen concentrations in each chamber. Here we define net flux as both background oxygen consumption and oxygen diffusion into the system. Flux at 100% air saturation was measured with fully oxygenated Ringer’s media. To measure net flux at lower oxygen saturations, Ringer’s media was deoxygenated to the desired level with nitrogen gas. 85% air saturation was chosen because cardiac MO_2_ measurements over the six minutes typically deplete oxygen to approximately 92% of air saturation but do not exceed 85%. To determine net flux, oxygen concentration was measured in each chamber for 10 minutes and repeated in triplicate.

Biological repeatability between chambers was tested with yeast at 28°C. A cell suspension was prepared using 1 g of yeast per 10 ml of Ringer’s media supplemented with 5 mM glucose. In each chamber, 100 μl of the suspension was injected to account for variation in chamber volume. Oxygen consumption was measured for 10 minutes in triplicate. MO_2_ was calculated as above to confirm there were no differences among chambers.

In order to assess metabolic consistency over the time-course of the experiment as well as chamber repeatability, hearts from four fish were isolated, and glucose metabolism was assayed in each chamber at 28°C. Hearts were randomly assigned to one of the four chambers and cycled through each of them, with media exchange between each measurement. Three blank measurements were run at the conclusion of the experiment. MO_2_ was calculated as above and then regressed against relative time of initial oxygen measurement per cycle to determine metabolic rate consistency of heart tissue over time.

### Substrate Specific Metabolism

For teleost ventricles, substrate specific metabolism was measured under four conditions: 1) 5 mM glucose, 2) fatty acids (1mM Palmitic acid conjugated to fatty-acid free bovine serum albumin), 3) LKA: (5mM lactate, 5mM hydroxybutyrate, 5 mM ethyl acetoacetate, 0.1% ethanol) and 4) non-glycolytic endogenous metabolism (no substrate). With the exception of glucose measurements, glycolytic enzyme inhibitors (20 mM 2-deoxyglucose and 10 mM iodoacetate) were added to all substrates to inhibit residual glycolytic activity. These substrate concentrations are commonly used for assaying substrate metabolism in teleost ventricles [19–22]. As outlined above, MO_2_ was measured over six minutes for each substrate. Measurements occurred at either 12°C or 28°C, the temperature at which the fish had been acclimated. After measurement, hearts were placed in a media bath of the next substrate while media was exchanged in each chamber. Substrate metabolism was measured in the above order for each individual heart (Fig 1C).

### Fatty acid saponification

Palmitic acid was conjugated to fatty acid-free bovine serum albumin (BSA) in order to facilitate cellular uptake by the heart ventricles. A solution of 4 mM sodium palmitate and 150 mM NaCl was prepared and heated to 70°C until fully dissolved. A second solution of 0.68 mM fatty-acid free BSA and 150 mM NaCl was prepared, sterile filtered, and warmed to 37 °C. Once dissolved, equal parts hot palmitate solution was transferred in small increments to the BSA solution, while stirring. The final solution was stirred for 1 hour at 37°C and stored at −20°C until use. Conjugation produces a final concentration of 2 mM palmitate: 0.34 mM BSA (6:1 FA:BSA), which is at a 2X concentration of the final working solution used in metabolic measurements.

### Statistical analysis

Raw data processing and statistical analyses were conducted using RStudio 1.1.463.

## Results

The HTMR is a simple custom design composed of a plexiglass water bath enclosing micro-respiration chambers connected to a multi-channel oxygen meter (Fig 1A). For a 10-chamber system, the approximate cost per chamber is $1870, including the cost of the oxygen meter and stir-plate. The full cost of the system is broken down in Table 1. The oxygen meter itself represents the highest cost (~$14,000 for ten inputs). followed by 1 ml glass chambers with lid containing two injection ports (~$300 each). Optical-fiber cables and sensor spots combined are approximately $95 each. A multi-place stir-plate is also necessary (~$900).

**Table 1.** Cost breakdown of HTMR system.

| Description | Manufacturer | Product # | Cost |
| --- | --- | --- | --- |
| 10-channel microfiber-optic oxygen meter | PreSens | 300000008 | \$13,966 |
| O<sub>2</sub> Sensor Spots | PreSens | 200001526 | \$26 |
| Fiber optical cable | PreSens | 200001731 | \$69 |
| Glass chamber | Unisense | MR-CH1 | \$88 |
| Glass chamber lid with two microinjection ports | Unisense | MR-CH INJECT<br>LID | \$183 |

### Methodological validation

Net flux was tested at multiple oxygen concentrations. In the chambers, background activity was negligible (2.038 pmol*s^−1^), when flux was measured at 100% air saturation (Fig 2A). In contrast, the average oxygen consumption rate of *F. heteroclitus* ventricles with glucose is 39.818 pmol*s^−1^, thus flux at 100% air saturation is approximately 5%. Similarly, at 85% air saturation (Fig 2B), flux was negligible (−0.324 pmol*s^−1^), representing less than 1% of cardiac glucose MO_2_. In contrast, flux was somewhat high at 50% with an average flux −14.429 pmol*s^−1^, likely due to high leak (S1 Fig). However, for all three oxygen concentrations, flux was equal among chambers by one-way ANOVA testing (p=0.696, p=0.643, p=0.733 at 100%, 85% and 50% air saturation, respectively).

**Fig 2.**
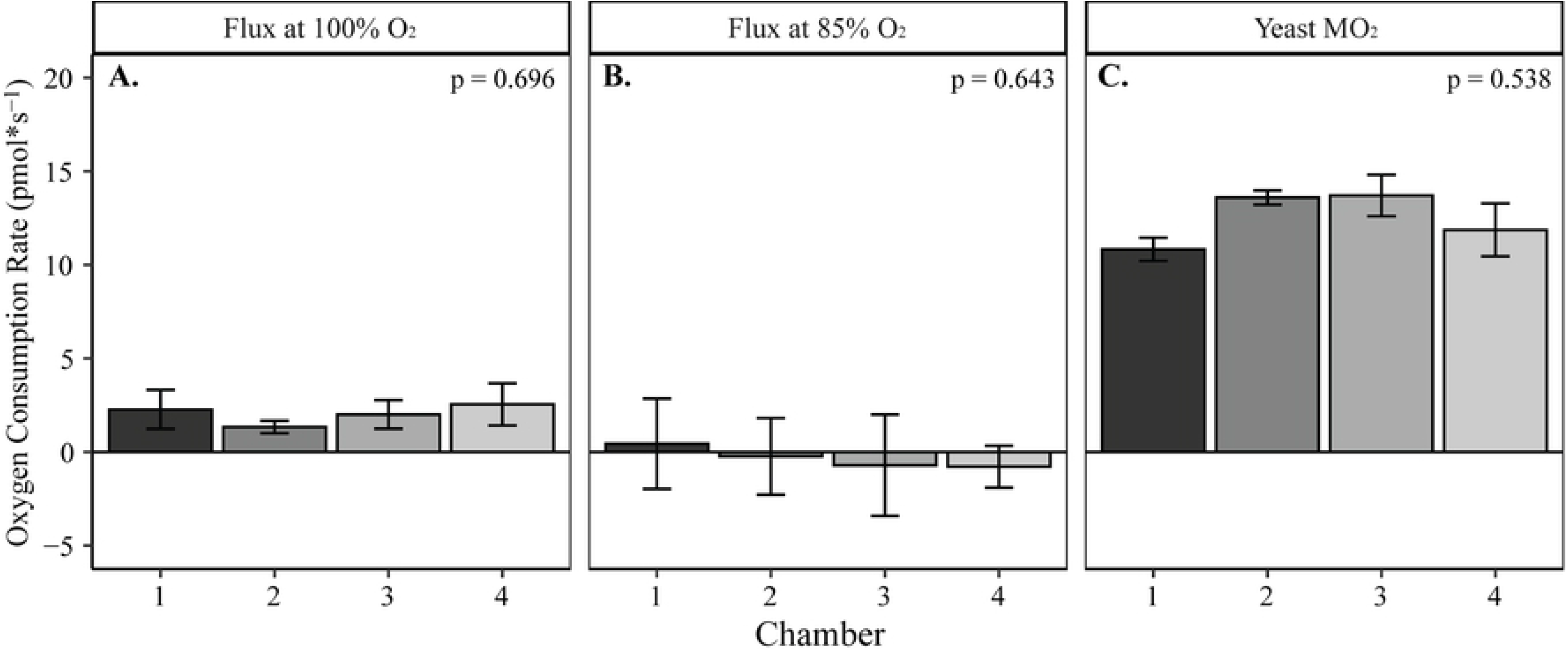
Chamber reproducibility in flux and metabolism. Flux was measured at 100% (A), 85% (B) in all four chambers of the HTMR. Additionally, MO_2_ of a standardized yeast cell suspension, was measured in each chamber (C). Oxygen consumption rates were calculated in pmol*s^−1^ and are represented by the mean with standard error bars (n=3 for each measurement). A one-way ANOVA was used for each measurement to test that there were no differences among chambers.

To test biological repeatability among the chambers, yeast metabolism per chamber was measured. Average MO_2_ was 12.502 ± 1.907 pmol*s^−1^, and there were no significant differences in metabolism between each of the chambers (ANOVA, p = 0.538; Fig 2C). This MO_2_ for yeast is approximately 40-fold higher than the net flux at 85% air saturation. In addition to yeast measurements, heart ventricles were measured across all four chambers over a 45-minute time period to validate both repeatability among chambers and that ventricles can maintain consistent metabolic activity over time. Among the four replicates, there was no significant difference in metabolic rate when regressed against time (linear model, p = 0.657; Fig 3A). Additionally, there were no significant differences in metabolic activity among the four chambers for each heart (ANOVA, p = 0.363; Fig 3B).

**Fig 3.**
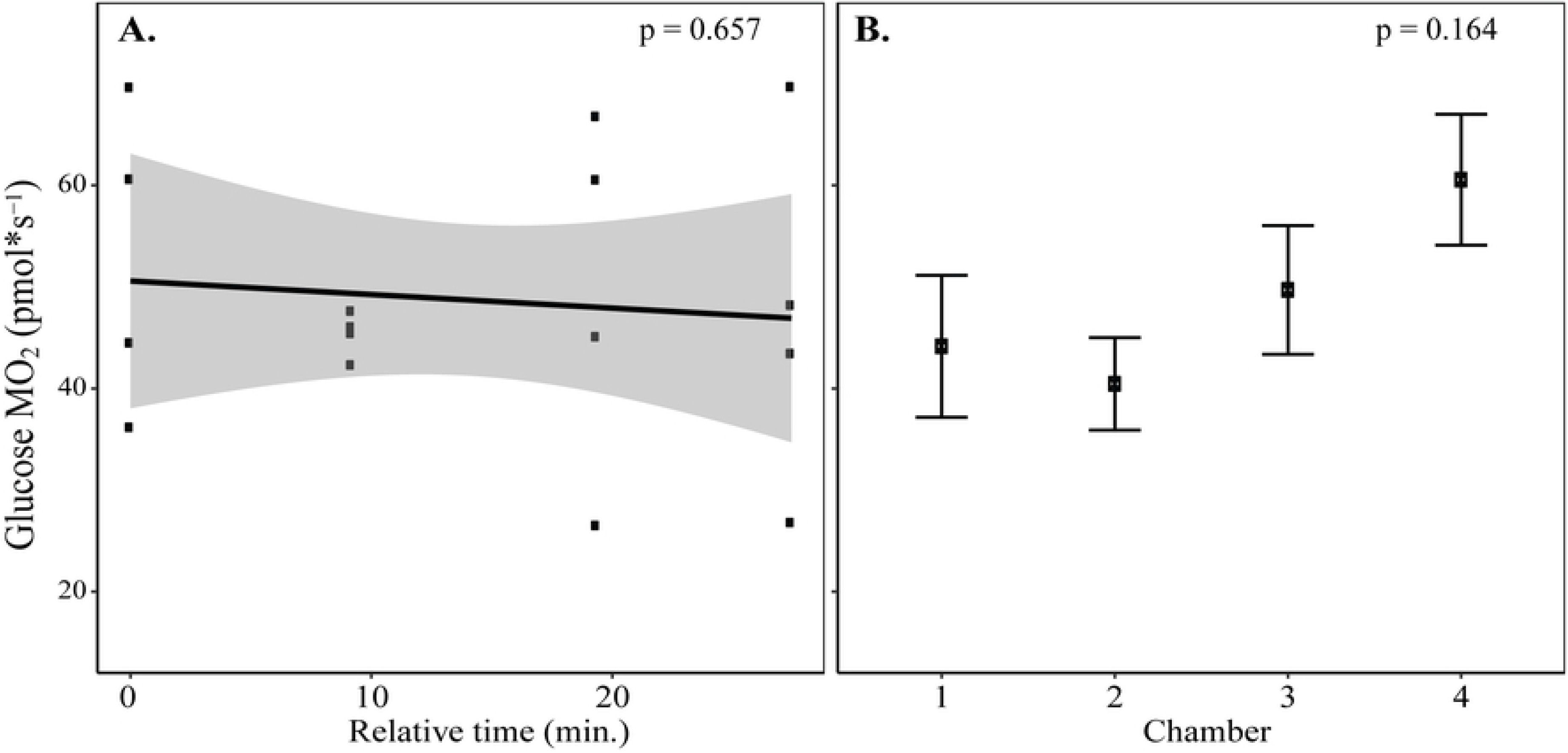
Cardiac glucose metabolism over time within the HTMR. *F. heteroclitus* ventricles (n=4) in Ringer’s with glucose media were rotated through the chambers and measured at 28°C. (A) Linear regression of cardiac glucose MO_2_ for the four replicated time periods (p=0.657) (B) Average metabolic rate per chamber, represented by the mean and standard error bars (ANOVA, p = 0.164).

### Substrate specific utilization

Heart ventricles were measured at corresponding acclimation temperatures; thus, temperature represents both the physiological effects of acclimation and the direct effect of temperature on MO_2_. All hearts were metabolically active under each of the four substrate conditions (Fig 4). For substrate specific metabolism, data was analyzed separately for individuals measured at 12°C or 28°C. There is a large inter-individual variation in substrate specific metabolism, which reduces the statistical power to reject the null hypothesis of no difference among substrates. To avoid this type II error, we apply paired t-tests that compare substrates within each individual and use Bonferroni’s test to correct for multiple tests [23]. Glucose, FA, and LKA metabolism were significantly greater than endogenous (p=0.001, Bonferroni’s corrected p=0.006); except FA at 12°C (p=0.03; Bonferroni’s corrected p=0.18; Fig 4A). Glucose metabolism was significantly greater than FA and LKA (Bonferroni’s corrected p=0.006) at both 12°C and 28°C. FA metabolism was significantly greater than LKA metabolism at 28°C (Bonferroni’s corrected p=0.006; Fig 4B) but not at 12°C (p=0.5; Fig 4A).

**Fig 4:**
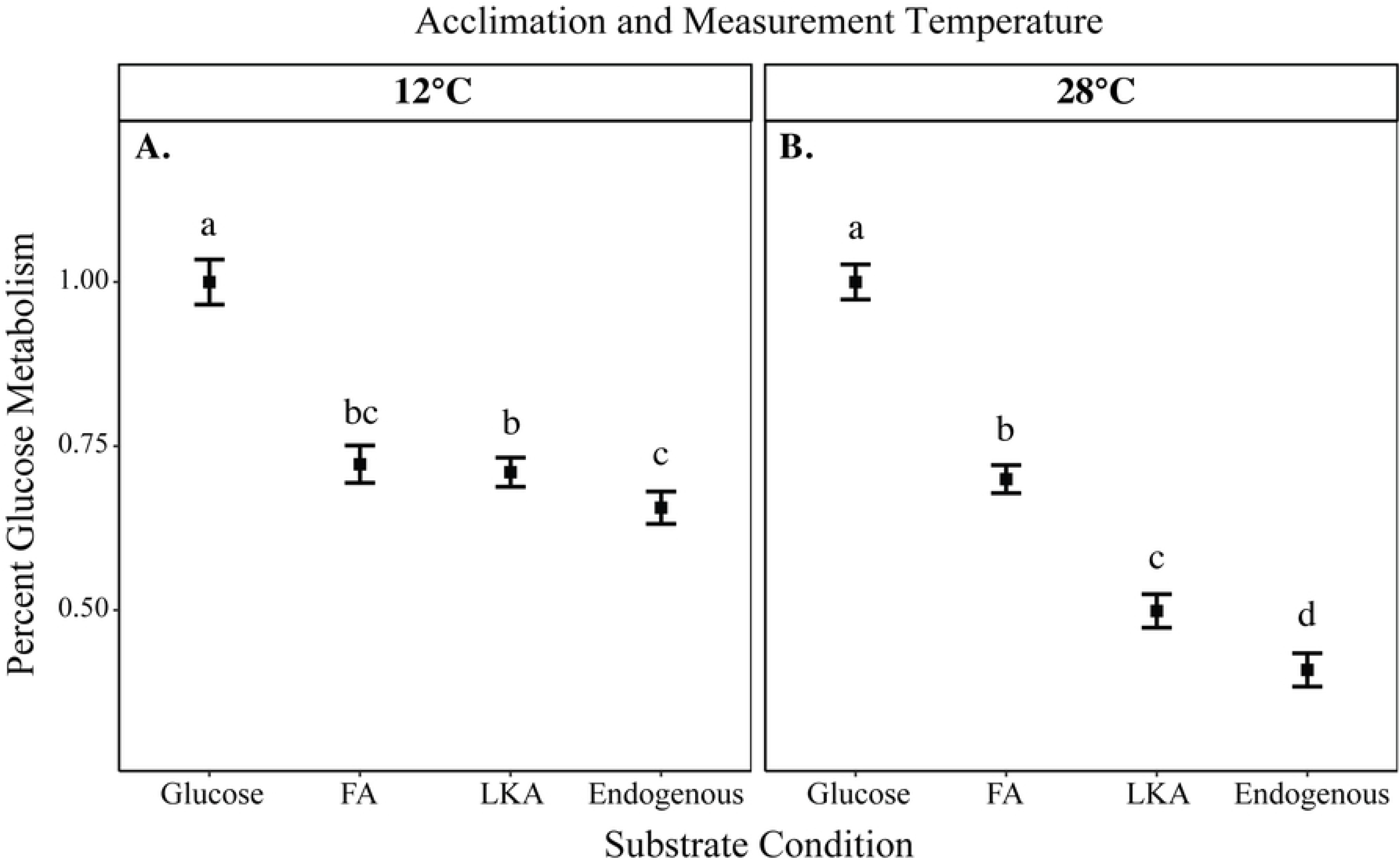
Heart ventricle substrate specific metabolism. Substrate specific ventricle MO_2_ as a percent of glucose consumption, represented by the mean with standard error bars. Percent glucose metabolism is relative to the mean glucose metabolism of all individuals measured at each temperature. Metabolism was measured at 12°C (A) and 28°C (B). A paired t-test was used to compare substrate metabolism within individuals, with Bonferroni test correction. Letters indicate significance levels between each substrate. At 12°C, n = 95. At 28°C, n = 105.

The log_10_ substrate specific MO_2_ from Maine and Massachusetts acclimated to 12°C and 28°C determined here can be compared to MO_2_ measured in Oleksiak *et. al* (2005) for Maine individuals acclimated to 20°C using an ANCOVA with log_10_ body mass and temperature as linear covariates. There were no significant differences (p = 0.55, 0.15, and 0.85 for glucose, FA and LKA respectively) and the least squares fall within 5% of one another.

### Allometric scaling of metabolism

Both body mass and heart ventricle mass were measured of each *F. heteroclitus* individual measured at each temperature. The mean body mass of individuals measured at 12°C and 28°C was 9.11 ± 2.87 g and 9.32 ± 2.90 g, respectively, and was not significantly different between temperatures. Additionally, average ventricle masses were 0.013 ± 0.005 and 0.010 ± 0.004 for 12°C and 28°C, respectively and did not significantly differ between acclimation temperatures. In *F. heteroclitus* body mass and heart mass are highly correlated (linear regression at 12°C R^2^= 0.74, p<0.0001; for 28°C, R^2^= 0.66, p < 0.001) thus, body mass was used to correct for variation due to mass between individuals, as done previously [24]. Body mass explained a significant amount of the variation (30-70%) in metabolism among individuals for all conditions (Fig 5). Variance explained by body mass (R^2^), was higher at 12°C than at 28°C (Fig 5, S1Table). For glucose MO_2_, allometric scaling was identical (to the 2^nd^ significant digit) to previous determinations and nearly the same as in Jayasundara *et al.* (2015). Examining the effect of temperature and substrates, allometric scaling coefficients (S1 Table), were between 0.65 to 1.29. While body mass contributed significantly to the variation between individuals, there was no effect of sex on cardiac metabolism by linear regression at each substrate-temperature combination. A three-way ANOVA including substrate, body mass and sex showed no significant differences between males and females in cardiac metabolism at 12°C or 28°C (p = 0.0963 and p= 0.4143, respectively).

**Fig 5.**
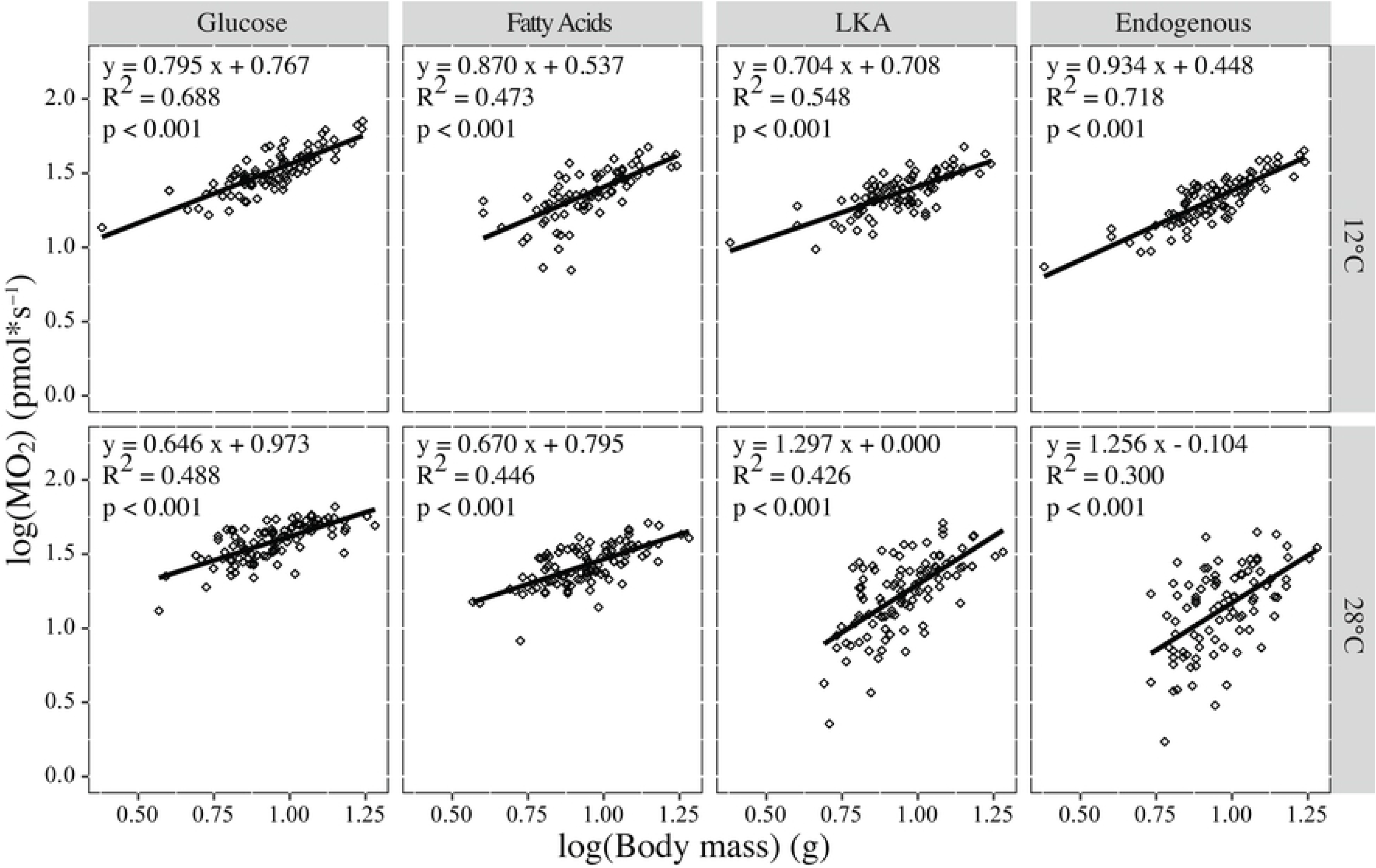
Allometric scaling relationship of substrate specific metabolism. Regression of log MO_2_ versus log body mass with each substrate. Top panels represent fish measured at 12°C while bottom panels represent 28°C. Hearts were measured as described in Fig 1 with glucose, fatty acids, LKA and endogenous metabolism. At 12°C, n= 105 and at 28°C n=95. For full data on regression slopes, see S1 Table.

## Discussion

### Design

The HTMR provides a simple custom design for measuring small biological samples that allows higher throughput measurements at lower costs. While cost is still not negligible, the system as described here, including the oxygen meter, sensors, stir control and chambers reduced cost by up to 30% compared to other manufacturers. The system costs approximately 10-fold less than the Agilent Seahorse and does not suffer from expensive disposable chambers and reagents. Using the 10 channel PreSens meter, combined with the custom chamber design, cost per chamber was approximately $1870 which was 70% and 80% of the cost compared to Unisense OPTO MicroOptode and Loligo OX11875 Witrox systems, respectively. The PreSens oxygen meter was primarily chosen due to the flexibility of the 10-channel system. Similar multi-channel systems such as the Strathkelvin Instrument use Clark electrodes, while PreSens uses smaller, more precise optical oxygen sensors. Unlike Clark electrodes, the optical sensors do not consume oxygen, are less expensive and have faster response times. Additionally, the fiber optical cables, and oxygen meter can be used with other sensor designs for applications such as whole animal metabolism [25]. This allows additional flexibility and is more economical, as the instrument can be used for measuring both tissue-specific, whole animal metabolism and numerous applications involved with oxygen measurement. Assuming a large intended sample size, with four substrate measurements per heart, this instrumentation design becomes more and more economical as you increase the number of chambers, making it useful for large phenotypic analyses.

Certain limitations were encountered while developing the multi-chamber system that are important to highlight. For the HTMR, several oxygen sensors were evaluated, but the sensor spots were chosen for their ease of use and durability. They are long lasting and customizable in their placement, allowing for repeated use. Other sensors, such as profiling oxygen microsensor probes were tested; however, they were more fragile and cumbersome to use. Temperature control is also essential for consistent and repeatable measurements. Temperature has a large impact on oxygen solubility; thus, precise temperature control is necessary and was closely regulated and monitored during measurements.

Chamber mixing was another important factor to control. Without thorough mixing from the stir-bars, oxygen measurements are inconsistent and inaccurate. This is an advantage of this system design over other multi-well plate style oxygen readers in which the media is unstirred during measurement, but instead rely on mixing prior to measurement. Continuous stirring allows for longer measurement periods. The size of the mesh that holds the tissue above the stir bar was also optimized: very small mesh inhibits mixing, but too large mesh would not separate the tissue from the stir bar in the bottom of the chamber. Additionally, nylon mesh was used over steel mesh, as it did not as readily retain air bubbles.

Finally, leak was tested extensively. At 85% air saturation, background flux (O_2_ use not associated with biological sample) was small (< 1 pmol*s^−1^) compared to heart ventricle and yeast MO_2_. To account for any amount of flux, blank measurements were taken throughout runs and corrected for in each chamber, with no significant differences in leak among chambers.

Initially, the Unisense microinjection lids were chosen for flexibility; however, after completing tests and measurements, the manufacturers released information that this particular model was less airtight than other models. For future design construction, we recommend that researchers use single-port lids with a sufficient path length to further minimize leak through diffusion.

### Methodological Validation

The HTMR is sensitive to both substrates used to fuel heart ventricle metabolism and body mass. HTMR determinations were very similar to previously published data (within 5%) [19]. Additionally, substrate specific patterns, with highest rates supported by glucose, agree with previous measurements in *F. heteroclitus* [19]. Metabolism was unaffected by the time course for measuring the four substrate conditions. Ventricles continue to contract over the duration of the experiment and show no significant decline in metabolic activity (Fig 2A.). Importantly, body mass accounted for a significant amount of variation in these individuals following an allometric scaling pattern, and the log mass against logMO_2_ linear regression has nearly identical slopes to those determined by others [15,19]. These data suggest that the system is both precise because the variation among samples did not obscure substrate or body mass effects and accurate in that substrate specific metabolic rates are similar to previous measures [19] and have allometric scaling coefficients very similar to published data [15].

## Conclusions

The HTMR was designed to measure metabolism in many individuals, because of the large individual variation and adaptive significance of the trait. We were specifically interested in cardiac metabolism in *F. heteroclitus*, as this species shows large inter-individual variance in cardiac metabolism and the mRNAs associated with this variance [19], and these patterns may hold true for many species. To better understand the physiological and evolutionary importance of this variance requires many individuals which the HTMR allows. For example, in this study, metabolism was quantified in approximately 200 ventricles in only 10 days. Although here the system is tested only with *F. heteroclitus*, this system could easily be extended to study other types of tissue-specific metabolism in many individuals. The decreased cost and efficiency of the design can have countless applications, allowing for high-throughput measurement of tissue metabolism that can enhance our understanding of the adaptive importance of these traits.

## Acknowledgements

The authors would like to thank Moritz A. Ehrlich for assistance with data collection in the substrate metabolism experiments.

## Supporting information

**S1 Fig.**
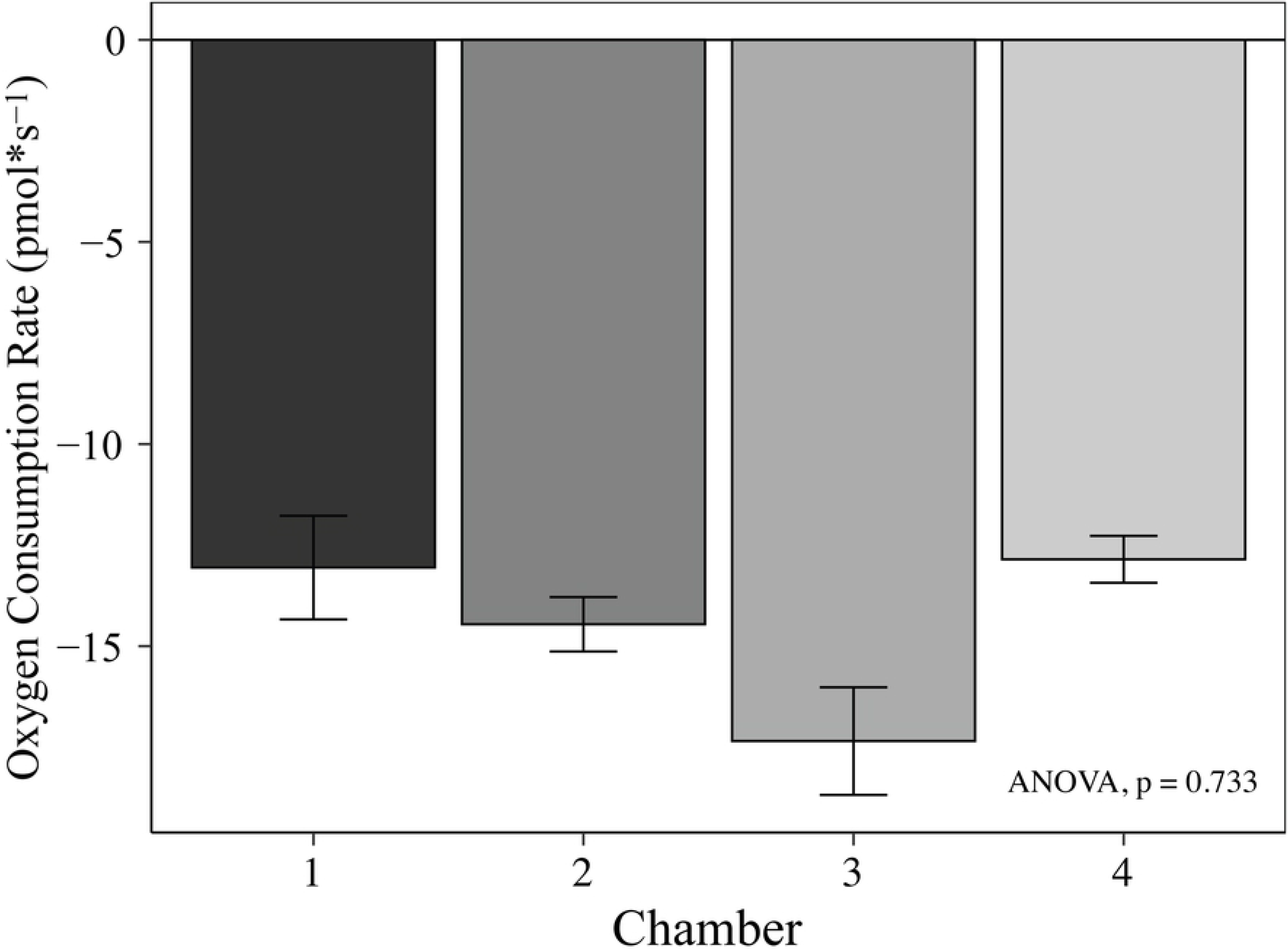
Chamber reproducibility in flux at 50% air saturation. Flux was measured at 50% air saturation in all four chambers of the HTMR. Oxygen consumption rates were calculated in pmol*s^−1^ and are represented by the mean with standard error bars (n=3 for each measurement). A one-way ANOVA was used to test variance between chambers (p=0.733).

**S2 Fig.**
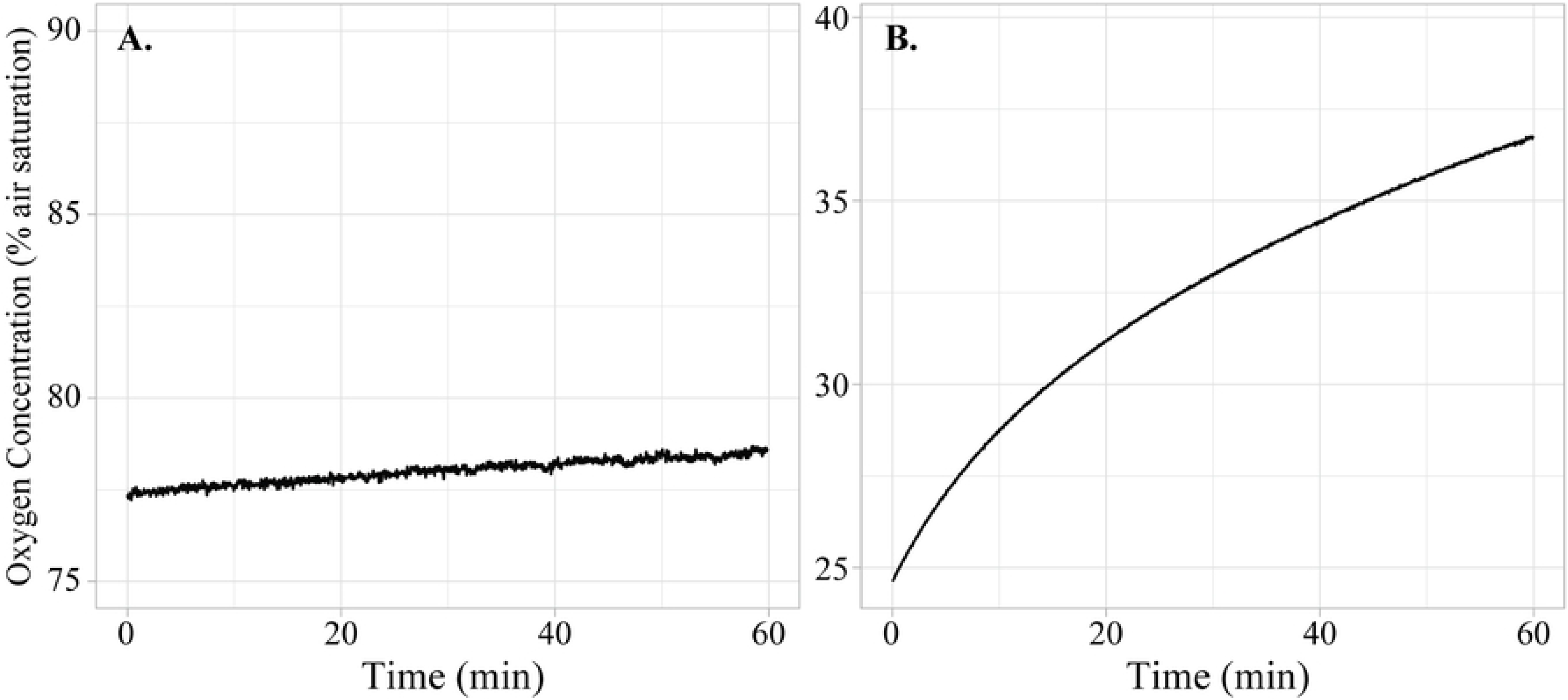
Chamber Leak over Time. Leak was tested at both high (A) and low (B) oxygen concentrations. Note the y-axes are different values but are incrementally scaled the same.

**S1 Table.** Descriptive statistics of body mass regression analysis. Slope and y-intercept data of the allometric relationship of body mass with substrate specific metabolism derived from a linear model of the log of metabolic rate against the log of body mass per substrate by temperature.

|  | 12°C |  |  |  | 28°C |  |  |  |
| --- | --- | --- | --- | --- | --- | --- | --- | --- |
|  | Glucose | FA | LKA | Endogenous | Glucose | FA | LKA | Endogenous |
| **Slope (b)** | 0.7951 | 0.8698 | 0.7042 | 0.9339 | 0.6462 | 0.6702 | 1.2967 | 1.276 |
| **y-intercept (a)** | 0.7666 | 0.5374 | 0.7076 | 0.4484 | 0.9734 | 0.7955 | -0.0004 | -0.1046 |
| **R<sup>2</sup>** | 0.6882 | 0.4726 | 0.5478 | 0.7184 | 0.4876 | 0.4464 | 0.2995 | 0.4259 |
| **N** | 94 | 80 | 91 | 90 | 101 | 99 | 101 | 89 |
| **p-value** |  |  |  |  |  |  |  |  |
| **Slope (b)** | 5.19E-25 | 1.89E-12 | 5.17E-17 | 6.04E-26 | 4.78E-16 | 4.21E-14 | 1.42E-13 | 2.86E-08 |
| **y-intercept (a)** | 1.65E-25 | 7.13E-07 | 2.19E-18 | 3.00E-11 | 6.48E-28 | 6.69E-19 | 0.998 | 0.609 |
| **Standard Error** |  |  |  |  |  |  |  |  |
| **Slope (b)** | 0.0558 | 0.1041 | 0.0678 | 0.0623 | 0.0666 | 0.0758 | 0.1513 | 0.2092 |
| **y-intercept (a)** | 0.0528 | 0.0996 | 0.064 | 0.059 | 0.0634 | 0.0719 | 0.1452 | 0.2038 |
| **95% Confidence Interval** |  |  |  |  |  |  |  |  |
| **Slope (b)** | 0.6835 to 0.9067 | 0.6617 to 1.0779 | 0.5686 to 0.8399 | 0.8092 to 1.0585 | 0.5130 to 0.7793 | 0.5187 to 0.8218 | 0.9941 to 1.5993 | 0.8575 to 1.6944 |
| **y-intercept (a)** | 0.6610 to 0.8723 | 0.3382 to 0.7366 | 0.5796 to 0.8356 | 0.3304 to 0.5664 | 0.8465 to 1.1003 | 0.6518 to 0.9392 | -0.2908 to 0.2901 | -0.5123 to 0.3030 |
| **Significance Between Slopes\*** | Endogenous | NS | Endogenous | Glucose LKA | LKA Endogenous | LKA Endogenous | Glucose FA | Glucose FA |

\* Comparisons are within a single temperature between substrates based upon 95% confidence intervals presented above.

## References

1. Crawford DL, Schulte PM, Whitehead A, Oleksiak MF. Evolutionary physiology and genomics in the highly adaptable killifish (Fundulus heteroclitus). Compr Physiol. 2020;In Press.

2. Wilson AC, Carlson SS, White TJ. Biochemical Evolution. Annu Rev Biochem. 1977;46: 573–639. Available: www.annualreviews.org

3. Pierce VA, Crawford DL. Phylogenetic analysis of thermal acclimation of the glycolytic enzymes in the genus Fundulus. Physiol Zool. 1997/11/15. 1997;70: 597–609. doi:10.1086/515879

4. Bacigalupe LD, Nespolo RF, Bustamante DM, Bozinovic F. The quantitative genetics of sustained energy budget in a wild mouse. Evolution (N Y). 2004/04/08. 2004;58: 421–429. doi:10.1111/j.0014-3820.2004.tb01657.x

5. Brown JH, Gillooly JF, Allen AP, Savage VM, West GB. Toward a metabolic theory of ecology. Ecology. 2004. doi:10.1890/03-9000

6. Das J. The role of mitochondrial respiration in physiological and evolutionary adaptation. BioEssays. 2006;28: 890–901. doi:10.1002/bies.20463

7. Crawford DL, Oleksiak MF. The biological importance of measuring individual variation. J Exp Biol. 2007/04/24. 2007;210: 1613–1621. doi:10.1242/jeb.005454

8. Rønning B, Jensen H, Moe B, Bech C. Basal metabolic rate: Heritability and genetic correlations with morphological traits in the zebra finch. J Evol Biol. 2007/08/24. 2007;20: 1815–1822. doi:10.1111/j.1420-9101.2007.01384.x

9. Galtier N, Jobson RW, Nabholz B, Glémin S, Blier PU. Mitochondrial whims: Metabolic rate, longevity and the rate of molecular evolution. Biology Letters. Royal Society; 2009. pp. 413–416. doi:10.1098/rsbl.2008.0662

10. Nilsson JÅ, Åkesson M, Nilsson JF. Heritability of resting metabolic rate in a wild population of blue tits. J Evol Biol. 2009/08/18. 2009;22: 1867–1874. doi:10.1111/j.1420-9101.2009.01798.x

11. Pettersen AK, Marshall DJ, White CR. Understanding variation in metabolic rate. J Exp Biol. 2018/01/13. 2018;221. doi:10.1242/jeb.166876

12. Itazawa Y, Oikawa S. A quantitative interpretation of the metabolism-size relationship in animals. Experientia. 1986;42: 152–153. doi:10.1007/BF01952441

13. Itazawa Y, Oikawa S. Metabolic rates in excised tissues of carp. Experientia. 1983;39: 160–161. doi:10.1007/BF01958874

14. Burggren W, McMahon B, Powers D. Environmental and metabolic animal physiology. Environmental and metabolic animal physiology. 1991. Available: https://books.google.com/books?hl=en&lr=&id=7fQvbFlQBaQC&oi=fnd&pg=PA3&dq=comparative+animal+physiology,+environmental+and+metabolic+animal+physiology&ots=r9_H809WWx&sig=LbQEnkf5RZ2r5RmbsULLLvmeOl4#v=onepage&q=comparativeanimalphysiology%2Cenvironmenta

15. Jayasundara N, Kozal JS, Arnold MC, Chan SSL, Di Giulio RT. High-throughput tissue bioenergetics analysis reveals identical metabolic allometric scaling for teleost hearts and whole organisms. Zhang J, editor. PLoS One. 2015;10: e0137710. doi:10.1371/journal.pone.0137710

16. Tsuboi M, Husby A, Kotrschal A, Hayward A, Buechel SD, Zidar J, et al. Comparative support for the expensive tissue hypothesis: Big brains are correlated with smaller gut and greater parental investment in Lake Tanganyika cichlids. Evolution (N Y). 2015;69: 190–200. doi:10.1111/evo.12556

17. Sukhum K V., Freiler MK, Wang R, Carlson BA. The costs of a big brain: Extreme encephalization results in higher energetic demand and reduced hypoxia tolerance in weakly electric african fishes. Proc R Soc B Biol Sci. 2016;283. doi:10.1098/rspb.2016.2157

18. Warren DL, Iglesias TL. No evidence for the “expensive-tissue hypothesis” from an intraspecific study in a highly variable species. J Evol Biol. 2012;25: 1226–1231. doi:10.1111/j.1420-9101.2012.02503.x

19. Oleksiak MF, Roach JL, Crawford DL. Natural variation in cardiac metabolism and gene expression in Fundulus heteroclitus. Nat Genet. 2004/11/30. 2005;37: 67–72. doi:10.1038/ng1483

20. Podrabsky JE, Javillonar C, Hand SC, Crawford DL. Intraspecific variation in aerobic metabolism and glycolytic enzyme expression in heart ventricles. Am J Physiol Regul Integr Comp Physiol. 2000/11/18. 2000;279: R2344–8. doi:10.1152/ajpregu.2000.279.6.R2344

21. Sidell BD, Crockett EL, Driedzic WR. Antarctic fish tissues preferentially catabolize monoenoic fatty acids. J Exp Zool. 1995;271: 73–81. doi:10.1002/jez.1402710202

22. Sidell BD, Stowe DB, Hansen CA. Carbohydrate Is the Preferred Metabolic Fuel of the Hagfish (Myxine glutinosa) Heart. Physiol Zool. 1984;57: 266–273. doi:10.1086/physzool.57.2.30163712

23. Sokal RR, Rohlf FJ. Biometry. 4th ed. W. H. Freeman and Co.; 1969.

24. Pierce VA, Crawford DL. Variation in the glycolytic pathway: The role of evolutionary and physiological processes. Physiol Zool. 1996;69: 489–508. doi:10.1086/physzool.69.3.30164212

25. Drown MK, DeLiberto AN, Crawford DL, Oleksiak MF. An Innovative Setup for High-Throughput Respirometry of Small Aquatic Animals. bioRxiv. 2020; 2020.01.20.912469. doi:10.1101/2020.01.20.912469

